# Visual training induced occipital fast sleep spindle clustering in humans revealed by full-night HD-EEG recordings

**DOI:** 10.1101/2024.11.12.623185

**Authors:** Patrícia Gerván, Gábor Bocskai, Andrea Berencsi, Ferenc Gombos, Ilona Kovács

## Abstract

This study investigates the impact of extensive visual procedural training on the temporal organisation of sleep spindles in healthy young adults. We selected 39 participants aged 16-20 and employed high-density electroencephalography to assess spindle characteristics during two full nights of sleep, with daytime practising in a contour integration task in between the two nights. We utilised linear mixed models to comprehensively analyse the effects of age and training on basic, clustering- and rhythmicity-related spindle parameters. Our findings indicate no significant age effects in this age-range, and no significant change between the two nights with respect to slow spindles. Fast spindles demonstrated a significant increase in density after training, and we observed significant changes in spindle clustering and rhythmicity parameters as well. Local spindle density, train density, and the ratio of clustered spindles have increased, and inter-train interval decreased by the second night. These results contribute to the growing literature on sleep-dependent memory consolidation by demonstrating that spindle reorganisation occurs not only in motor tasks but also in visual learning contexts. The absence of age-related differences further highlights the robustness of these mechanisms across developmental stages. Our study emphasises the importance of spindle dynamics in procedural learning and suggests promising possibilities for future research into the neurophysiological basis of memory consolidation. By revealing the relationship between training and sleep spindle characteristics, our findings provide valuable insights into how sleep supports learning and memory processes in young adults, potentially informing interventions aimed at enhancing memory performance through sleep-related strategies.

## Introduction

Recent research on the mechanisms of sleep-dependent memory consolidation has increasingly focused on the role of sleep spindles (Fernandez and Lüthi, 2020; Niethard et al., 2018; Petzka et al., 2022; Peyrache and Seibt, 2020). Sleep spindles are cortical oscillatory patterns, ranging from 11 to 15 Hz, occurring during non-rapid eye movement (NREM) sleep, and are generated by the thalamic reticular nucleus. In addition to their established role in declarative and emotional memory formation (Diekelmann and Born, 2010), there is growing evidence of their involvement in procedural learning, particularly in the consolidation of motor memory (Boutin and Doyon, 2020). Current models suggest that following an initial acquisition phase, sleep spindles support the offline consolidation of memories, selectively reactivating learning-specific brain regions (Boutin and Doyon, 2020; Petzka et al., 2022). The temporal structuring of sleep spindles into so-called spindle trains has been highlighted as essential for motor memory formation (Boutin et al., 2024; Boutin and Doyon, 2020), although these studies primarily address the relationship between spindle organisation and post-sleep performance improvement, not specifically the reorganisation of spindles after practising. Notably, while evidence linking daytime practice to subsequent local spindle density change has been demonstrated in a small group of epileptic patients with implanted electrodes (Johnson et al., 2012), direct evidence of post-training spindle organisation and its broader generalisation to other forms of procedural learning remains lacking.

In this study, we investigate how extensive training in a procedural task impacts the temporal organisation of sleep spindles within low-level, learning-related brain regions. To address this, we compare key parameters of sleep spindle clustering before and after the behavioural training period using full-night, high-density electroencephalography (HD-EEG). The daytime training involved multiple repetitions of a visual contour integration task (Kovács and Julesz, 1994; Kozma et al., 2006) known to be a late-developing ability (Kovács et al., 1999), which improves in a sleep-dependent manner (Gerván and Kovács, 2007) and depends on occipital cortical areas (Gilad et al., 2013; He et al., 2024). Comparing the occipital NREM activity from nights before and after training, we observed an increase in local fast spindle density by the second night, alongside other clustering parameters that reflect the temporal organisation of sleep spindles following extensive visual training.

## Methods

We selected 39, 16-20-year-old participants from our already existing database (Gombos et al., 2022) where the relevant basic sleep spindle parameters and their topographical organisation does not seem to change across these age groups (Bocskai et al., 2023; Gombos et al., 2022). Participants were recruited via social media. Only healthy individuals without sleep or neurological conditions were included in the original sample of 39 participants. After data analysis, one 16-year-old male participant was excluded due to his extreme outlier values. Our final sample had 38 participants in two age groups: 16-year-olds (n = 18, mean age = 15.97 ± SD 0.44 years, f: 10) and 20-year-olds (n = 20, mean age = 21.29 ± SD 0.51 years, f:10). Each participant received a voucher worth HUF 20,000 (approx. EUR 50.00) for their involvement. The study was approved by the Ethical Committee of Pázmány Péter Catholic University (PPCU) for Psychological Experiments, and written informed consent was obtained from participants and/or their parents.

Participants spent three consecutive full nights in the sleep laboratory of PPCU Budapest, where 128-channel HD-EEG polysomnography was recorded. They were asked to maintain their usual sleep-wake cycles for five days prior to the study, to avoid taking any drugs or caffeine on the day of the study, and to refrain from napping on study days. Participants went to bed between 10 p.m. and 11:30 p.m. and woke up spontaneously in the morning. For details on EEG recording procedures, hardware environment, data processing, and sleep spindle extraction methods, see Gombos et al. (2022). In this study, we excluded data from the adaptation night, and focused on data from 17 occipital electrodes (POOz, Oz, Iz, I1, OI1, POO3, I2, OI2, POO4, POO9h, POO11h, POO10h, POO12h, POO7, POO8, PO11, PO12) during the following two nights (night 1: N1, and night 2: N2).

Between N1 and N2, participants practised a contour integration task (Kovács and Julesz, 1994; Kovács et al., 1999; Kozma et al., 2006; Gerván and Kovács, 2007) at 8 a.m., 1 p.m., and 6 p.m., for approximately 15-20 minutes within each session (see Fig 1a). Stimuli were presented on a laptop computer screen. The images consisted of collinear chains of Gabor elements forming a horizontally positioned egg shape (target) against a background of randomly positioned and oriented Gabor patches (noise). The carrier spatial frequency of the Gabor patches was 5 c/deg, and their contrast was 95%. The spacing between the contour elements was kept constant (8λ, where λ is the wavelength of the Gabor stimulus), while the average spacing between the background elements was varied. The signal-to-noise ratio, defined by a D parameter (D = average background spacing/contour spacing), was varied between 1.2 and 0.5 in 15 steps. A new shape and background were generated for each stimulus, but all contours maintained the same general size and egg-like shape (see Fig. 1b; for more information on stimuli, see the OSF link below). In a two-alternative forced-choice (2AFC) procedure, participants were asked to indicate the direction in which the narrower part of the egg pointed. The stimulus was presented for 2000 milliseconds, followed by a fixation cross between stimuli (500 ms, or shorter if the participant responded more quickly). An adaptive staircase procedure ensured that each participant practised around their individual threshold level throughout each session consisting 180 trials.

**Figure 1.**
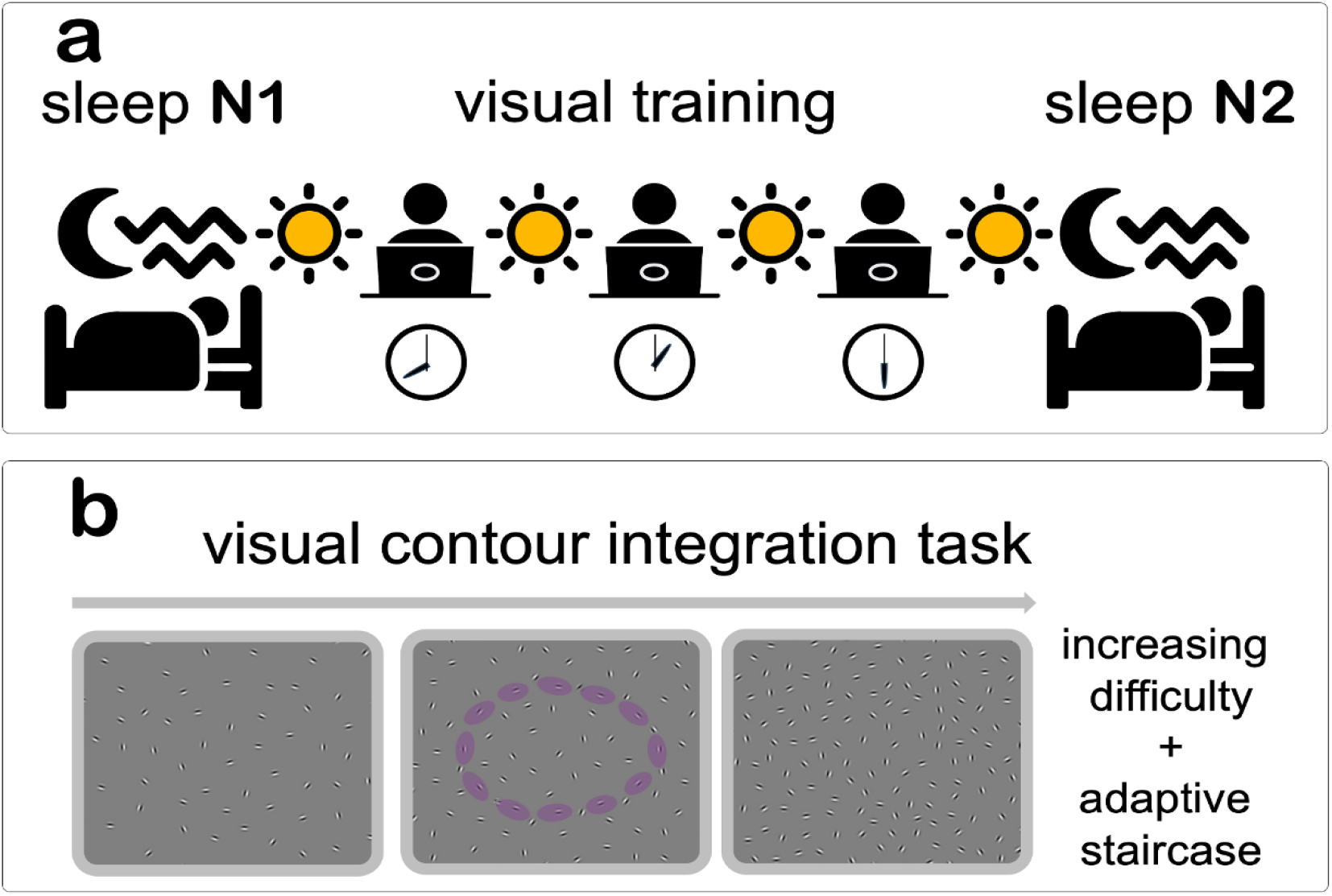
The visual training paradigm. **a**, Participants completed three practice sessions of a visual task during the daytime between two consecutive nights of full-night HD-EEG polysomnography recordings. **b**, The visual task involved contour integration, with an adaptive staircase method used to individually adjust task difficulty to threshold levels.

**Figure 2.**
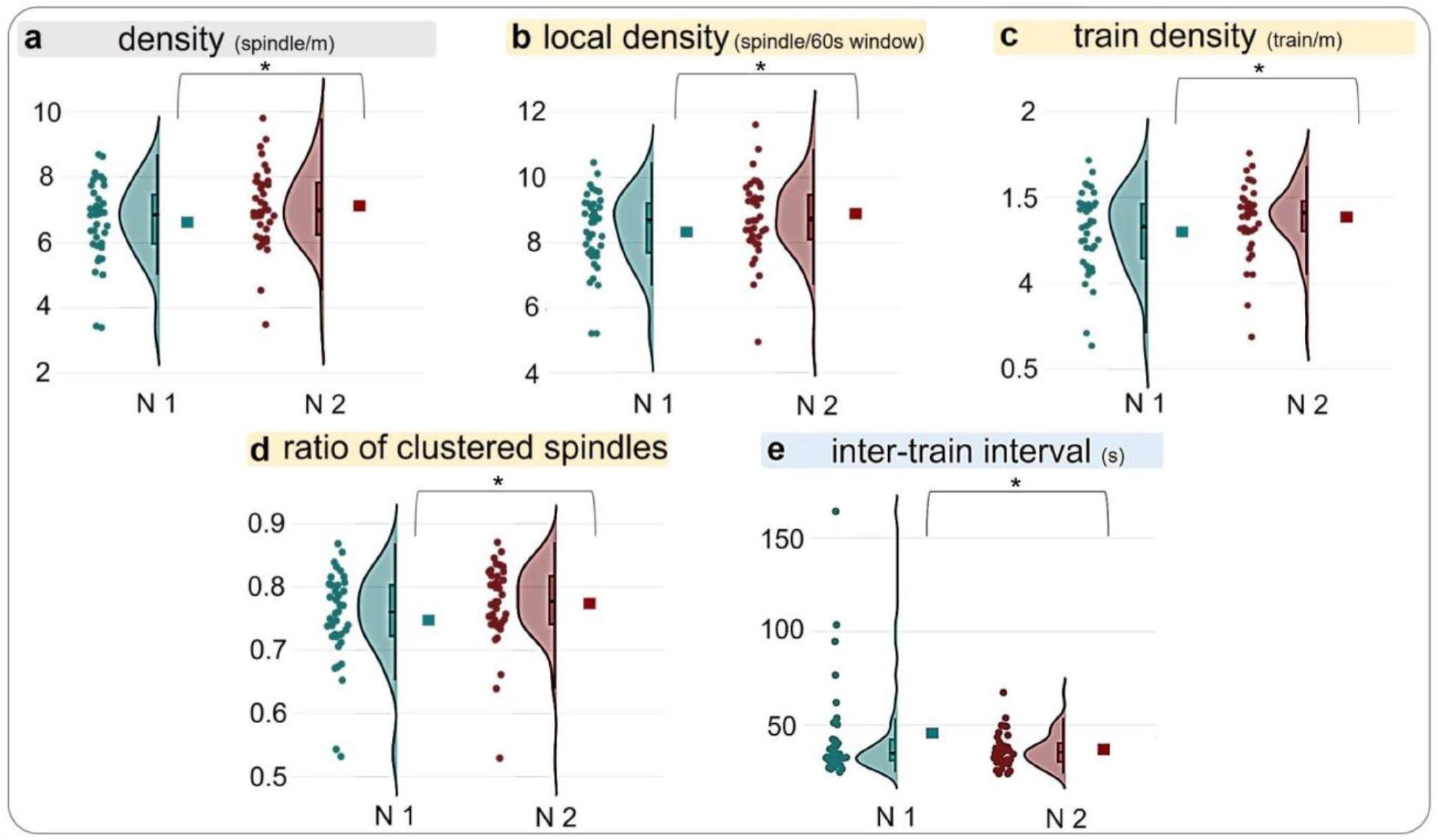
Raincloud plots of the raw sleep spindle data with estimates of LMM. Scatter plots represent values of individual data points. Raincloud shapes depict kernel density estimates of the data (probability density of the data at different points). Within raincloud plots, box plots specify the range in which the middle 50% of all data points are present and solid horizontal lines indicate the median. Square markers illustrate linear mix model estimates. The panels depict (a) spindle density (spindles per minute), (b) spindle local density (number of spindles within a 60-second sliding window centred around each spindle), (c) spindle train density (spindle trains per minute), (d) ratio of clustered spindles relative to all detected spindles and (e) inter-train intervals (seconds) parameters during Night 1 (N1) and Night 2 (N 2). Brackets and asterisks indicate significant Night fixed effects predicted by the LMM.

Here we focused on the changes induced by visual training on slow (9-12 Hz) and fast sleep spindle (12-16 Hz) characteristics and clustering. We evaluated basic spindle characteristics being density (spindles per minute), duration (s), maximum amplitude (µV), and peak frequency averaged across 17 occipital electrodes. We also applied additional metrics to explore whether spindles follow specific temporal reorganisation patterns (Boutin et al., 2024). We assessed clustering by calculating spindle local density, defined as the number of spindles within a 60-second sliding window centred around each spindle; counting spindle trains (with spindles spaced at least 6 seconds apart) and termed it spindle train density (number of trains per minute); calculating the mean number of spindles per train; and determining the ratio of clustered spindles relative to all detected spindles. Furthermore, we evaluated spindle rhythmicity determining inter-train interval (s); inter-spindle interval within trains (s); and the average duration of spindles (s) within and out of trains.

To examine whether the effect of training was associated with age, age group was included as a fixed effect in the first wave of Linear Mixed Model analysis. Mixed models were used to assess the effects of age group and training on twelve spindle-related parameters, accounting for the correlations among repeated measures taken from the same subjects across different nights. The models (see the summary in Supplementary Table S1) included Age Group, Night, and the Age Group × Night interaction as fixed effects, with one random effect (intercept per participant, capturing the variability in sleep parameters across participants).

Data analysis was performed with SPSS, version 27.0 (SPSS Inc., Chicago, IL, USA). Linear Mixed Models were generated using restricted maximum likelihood (REML) and heterogeneous Toeplitz covariance structure for repeated measurement. The p-values for fixed effects calculated using Satterthwaites approximations and standard errors have been calculated using the Wald method, significance level was set at p<.05.

## Results

With respect to slow spindles there was no significant Age Group or Night effect, and their interaction was not significant either (see Supplementary Table S2).

With respect to fast spindles, five models revealed significant Night fixed effect, however none of the linear mixed models revealed a significant main effect of Age Group, and the interaction between Night and Age Group was not significant either (see Supplementary Table S3). In the absence of a significant Age Group effect we modified the models by omitting Age Group (see the summary in Supplementary Table S4).

With respect to the basic parameters, Night had a significant effect only on spindle density (β_N1_=6.65(.02), β_N2_=7.05(0.2); p=.014). Regarding the clustering variables, Night had a significant effect on local spindle density (β _N1_=8.36 (1.08), β_N2_=8.75 (1.09); p=.025), train density (β_N1_=1.3 (.04) β_N2_=1.37 (.04); p=.023), and the ratio of clustered spindles (β_N1_= .75 (.01), β_N2_ =.77 (.01); p=.014). With respect to the rhythmicity parameters, we found a significant effect of Night in the inter-train interval (β_N1_=43.56 (4.03), β_N2_ =37.08 (2.17); p=.02). Results are summarised in Table 1 and Figure 1. The entire analysis can be found in Supplementary Table S5, and parameters with non-significant Night effect are depicted in Supplementary Figure S1).

**Table 1.**
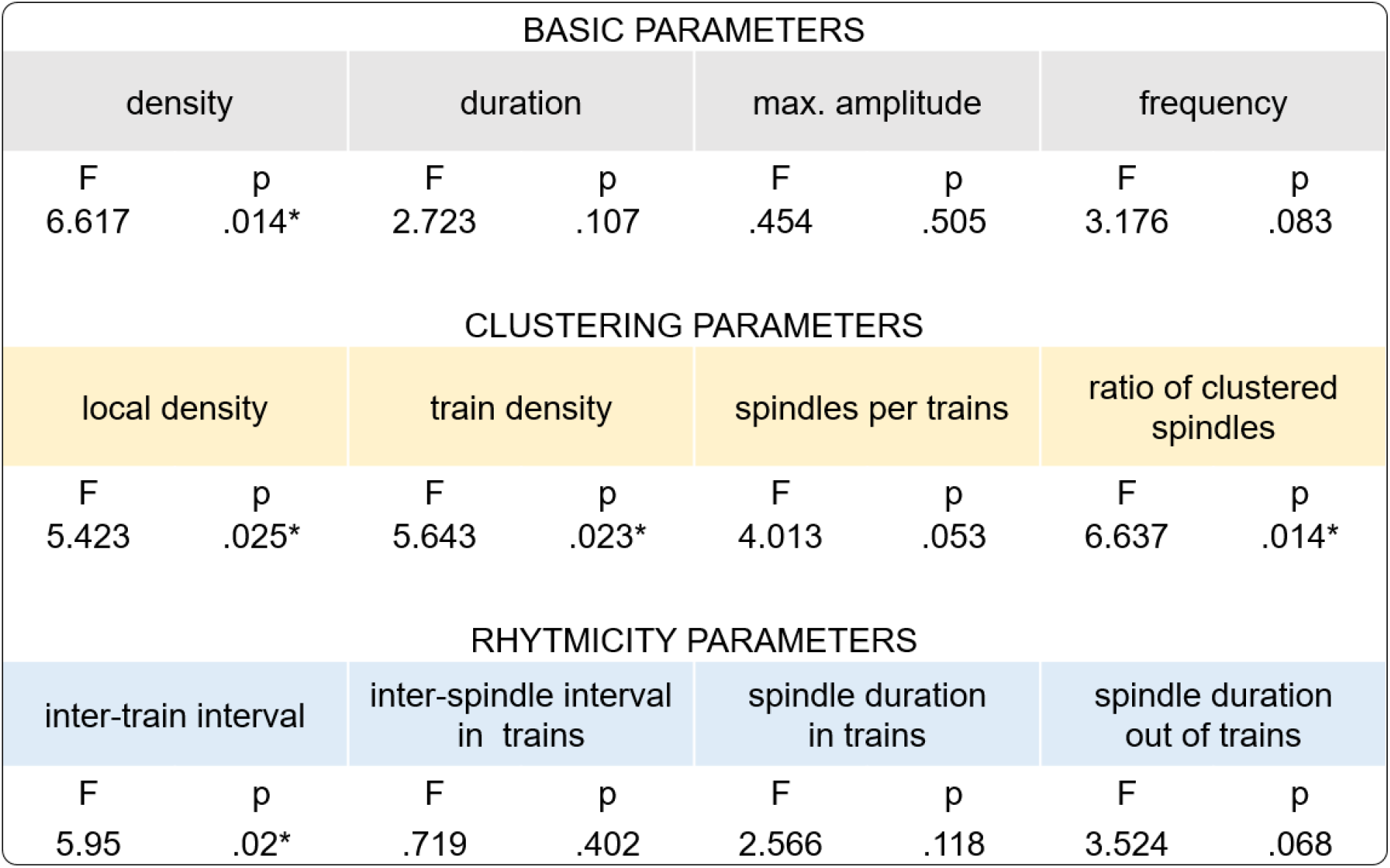
Night fixed effects of the linear mixed models for the analysed twelve fast spindle parameters.

## Discussion and conclusion

Our findings provide direct evidence that extensive procedural training reorganises the temporal structure of sleep spindles, contributing to the understanding of sleep-dependent memory consolidation mechanisms. While previous research has largely concentrated on the role of spindles in motor memory (Boutin and Doyon, 2020), our results extend this understanding to visual perceptual training. The significant increase in fast spindle density observed after training, coupled with alterations in clustering and rhythmicity parameters, suggests that spindles become increasingly organised into trains to facilitate memory consolidation in the occipital cortex.

These results align with current models proposing that spindle trains are integral to memory consolidation processes, selectively reactivating learning-specific brain regions during sleep (Boutin et al., 2024; Petzka et al., 2022). Importantly, our study highlights the absence of age-related differences in spindle dynamics in the given age-range, indicating that the reorganisation of spindles following training is robust across different developmental stages. This finding underscores the universality of spindle dynamics in procedural learning contexts.

Utilising high-density EEG allowed us to gather comprehensive evidence of post-training spindle reorganisation within a healthy population, extending earlier findings based on implanted electrodes (Johnson et al., 2012). The observed changes in spindle clustering parameters reinforce the notion that temporal structuring is crucial for procedural memory consolidation, supporting the hypothesis that spindle trains are essential for the effective consolidation of newly acquired skills.

In conclusion, our study emphasises the role of spindle dynamics in facilitating procedural learning and highlights promising directions for future research aimed at investigating the neurophysiological mechanisms underlying memory consolidation. By revealing the relationship between training and sleep spindle characteristics, we contribute valuable insights into the interplay between sleep and learning processes in young adults.

Supplementary information is available at the <u>OSF platform</u>.

## Acknowledgments

The authors thank the adolescents and young adults who participated in this project. This research was supported by the Hungarian National Research, Development and Innovation Office grants NK-104481 and K-134370 to I.K. It was made in the framework of the PPKE-BTK-KUT-23-1 project, with the support and funding provided by the Faculty of Humanities and Social Sciences of Pázmány Péter Catholic University.

